# Benefits and costs of contextual associations between real-world objects in the attentional blink

**DOI:** 10.64898/2026.08.06.743178

**Authors:** Lu-Chun Yeh, Daniel Kaiser

## Abstract

The attentional blink is a well-known phenomenon illustrating the limitations of human attention: When two visual targets are presented in rapid succession, identification of the second target is often impaired. While the attentional blink is known to attenuate when targets share perceptual features or category membership, real-world objects are also linked through contextual associations, shaped by objects typically occurring within the same environments. Here, we devised an attentional blink experiment in which we orthogonally manipulated contextual and categorical relationships between the two targets while controlling for their perceptual similarity. As the key result, contextual associations facilitated identification of the second target but impaired identification of the first target. These findings suggest that contextual associations yield distinct benefits and costs for visual cognition, where enhanced attentional access to subsequent targets is traded off against increased interference between targets.

## Introduction

Humans only have a limited capacity to process visual information. To adapt to a complex and cluttered visual environment, the visual system relies on attention to prioritize task-relevant information for further processing (Duncan, 1984). However, attentional resources are themselves limited (Luck & Ford, 1998). A well-known demonstration of these processing constraints is the attentional blink: when two targets are presented in rapid succession, identification of the second target is impaired if it appears shortly after the first, reflecting a temporal bottleneck in attentional processing (Weichselgartner & Sperling, 1987; Broadbent & Broadbent, 1987; Raymond et al., 1992).

Although the attentional blink has been robustly demonstrated in laboratory settings using simple stimuli such as letters and digits, humans appear surprisingly efficient at processing multiple events in rapid succession in natural environments, such as during cooking or driving in traffic. One reason may be that successive events in natural environments are often related to one another, allowing processing of one event to facilitate the next (Olivers, 2007). Consistent with this idea, previous studies using real-world objects have shown that the attentional blink is attenuated when the two targets share similar perceptual or categorical features (Kellie & Shapiro, 2004; Lindh et al., 2019).

However, real-world objects are related not only by perceptual and categorical similarity but also by contextual associations. Objects that frequently co-occur in the same scene are processed more efficiently, suggesting that contextual regularities facilitate object processing (Bar, 2004; Oliva & Torralba, 2007; Kaiser et al., 2019). These associations may be especially relevant for sequential behavior in natural settings, where one event often predicts the next. For example, after chopping vegetables, we typically place them into a cooking pot. Consistent with this idea, studies of visual search have shown that contextual associations guide attentional selection toward relevant objects (Torralba et al., 2006; Mack & Eckstein, 2011; Yeh et al., 2025). Although this evidence concerns spatial rather than temporal selection, it suggests that contextual associations can influence attentional prioritization more broadly. This raises the possibility that contextual associations between real-world objects may also alleviate the temporal bottleneck underlying the attentional blink. A potential mechanism for this is suggested by neuroimaging studies showing overlapping representations for contextually associated objects even when presented in isolation (Bar & Aminoff, 2003; Bonner & Epstein, 2021; Yeh et al., 2026). Such overlapping representations may yield a pre-activation of the representation of the second target in cases where the first target is contextually related (Lindh et al., 2019; Tang et al., 2022).

In the present study, we investigated how contextual associations influence target processing during the attentional blink. A key challenge is that objects sharing contextual associations often also share perceptual and categorical similarity. For example, spoons and knives frequently co-occur in kitchen contexts, but they are also visually similar and belong to the same semantic category (i.e., tools). This makes it important to disentangle the contributions of these three forms of object-relatedness. Moreover, previous research has shown that shared categorical representations between targets can attenuate the attentional blink (Lindh et al., 2019), raising the question of whether contextual and categorical associations differentially influence the attentional blink and whether their effects interact. To address these issues, we orthogonally manipulated the contextual and categorical relationships between the first (T1) and second (T2) targets while controlling for perceptual similarity. Our results showed that contextual associations facilitated detection of the T2 but impaired identification of the T1 during the attentional blink, suggesting that contextual associations yield distinctive benefits and costs for visual cognition.

## Methods

### Participants

Forty healthy volunteers (32 females, 8 males, age =25.43 ± 3.50 years) participated in the study. Sample size was determined to achieve approximately 80% power to detect a hypothetical medium effect (d = 0.5) at p < 0.05 (two-sided t-test), yielding a required sample size of N = 34. Six additional participants were recruited to allow counterbalancing across the 20 stimulus combinations (see below). All participants were native German speakers and had normal or corrected-to-normal vision. They provided written informed consent before participation and were compensated at a rate of 10 euros per hour. The study was approved by the Ethics Committee of Justus Liebig University Giessen and conducted in accordance with the 6th revision of the Declaration of Helsinki.

### Stimuli

We used a set of 24 real-world objects, which were also used in our previous neuroimaging studies (Yeh et al., 2025; Yeh et al., 2026). The stimuli stemmed from two contexts (12 garden objects and 12 kitchen objects) and two categories (12 non-tools and 12 tools). We organized the objects into six four-item sets, ensuring each set included one item from each condition (i.e., 1 garden tool, 1 garden non-tool, 1 kitchen tool, and 1kitchen non-tool). The objects within each set shared a similar overall shape (see Figure 1A). There were four exemplars per object, yielding 96 unique stimuli in total. Visual differences among objects within and between contextual and categorical conditions were examined using a VGG16 deep neural network as well as the HMAX hierarchical model of early visual processing. Neither model revealed significant differences when comparing objects within and between conditions. Detailed image-based analyses of visual similarity can be found in Yeh et al. (2026). Each participant was presented with three of the sets, yielding 20 possible combinations, which were fully counterbalanced across participants. A total of 204 images were generated as distractors. Specifically, the 96 original stimuli were first converted into texforms using a texture-synthesis algorithm (Long et al., 2018; Deza et al., 2019). Then, four texform images that did not share the same object identity were randomly selected and averaged to create each distractor image. All stimuli were placed on a white square background (5° × 5° visual angle).

**Figure 1.**
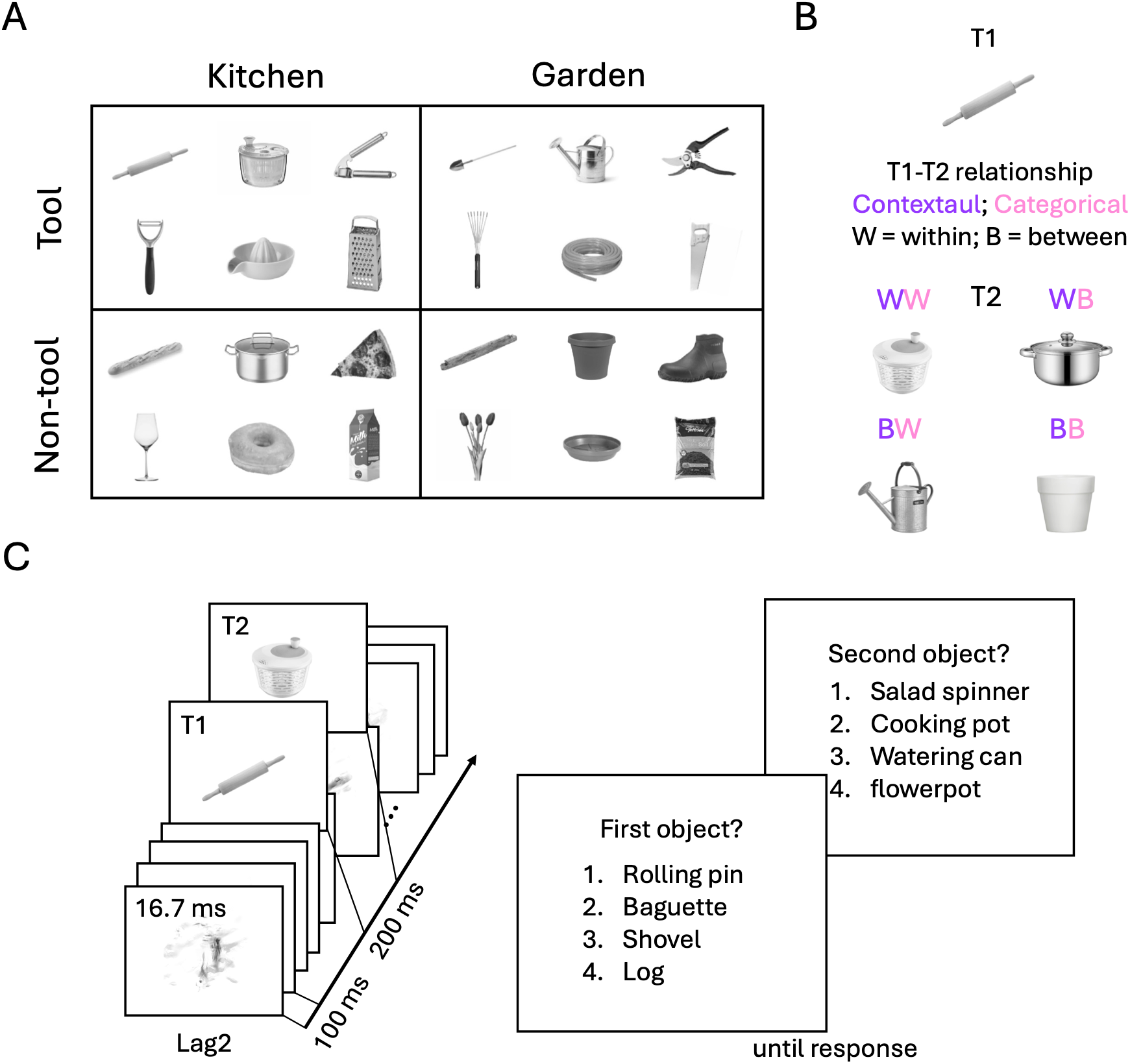
Stimulus, design, and task procedure. (A) Stimulus set. 24 objects were used in the experiments, selected from two contexts (kitchen and garden) and two categories (tool and non-tool), with objects shape-matched across the four sub-categories. (B) Illustration of T1-T2 pairings across conditions. Pairings vary by context (within vs. between) and category (within vs. between), yielding four combinations (WW, WB, BW, BB). (C) RSVP stream procedure. The four response options in the experiment were presented in German.

### Experimental Design and Paradigm

The attentional blink task employed a 2 (Lag: 2, 8) × 2 (Contextual relationship: between, within) × 2 (Categorical relationship: between, within) within-subjects factorial design. These factors were defined based on the relationship between the first target (T1) and the second target (T2) in a stream of images. Lag refers to the temporal interval between T1 and T2, with Lag 2 corresponding to 200 ms (with 1 distractor in between the targets) and Lag 8 to 800 ms (with 7 distractors in between). Contextual and categorical relationships were defined by whether the two targets belonged to different contexts or categories (between condition; e.g., one item from kitchen/tool and one from garden/non-tool) or the same context or category (within condition; e.g., both items from kitchen/tool or both from garden/non-tool). Each participant saw three object sets (12 objects). Each object was paired with the eight objects from the other sets, yielding 96 ordered pairs (including both A-B and B-A combinations). Objects were not paired within sets, and no pair shared overall shape, to minimize potential confounds from perceptual similarity. Each pair was presented twice at both Lag 2 and Lag 8, resulting in 384 trials in total and 48 trials per condition.

Each trial started with a fixation cross that appeared at the center of the screen for 500 ms to signal trial onset, followed by a stream of 19 images. Each image was presented for 16.7 ms with a stimulus-onset asynchrony (SOA) of 100 ms. Embedded into the stream of distracters, two targets (T1 and T2) were presented. T1 was always presented as item 5 in the stream, while T2 was either presented as item 7 (Lag 2) or item 13 (Lag 8). After the RSVP stream, participants reported the targets in order by selecting each target object from four response options (presented as German words), using the corresponding keys (digits 1 to 4). In addition to the target, the three distractors were objects from the same set, sharing the target’s overall shape. For example, when the target was a salad spinner (Salatschleuder), the other response options were a cooking pot (Kochtopf), a watering can (Gießkanne), and a flower pot (Blumentopf). The intertrial interval varied randomly between 800, 1000, and 1200 ms. The overall procedure is illustrated in Figure 1B.

### General Experimental Procedure

At the start of the experiment, participants filled out a questionnaire to check their familiarity with the 12 chosen objects. They were shown the German names for each item and asked if they recognized them. To guarantee proper identification during the subsequent task, the experimenter explained any objects that a participant did not know. After that, participants practiced a block of 12 trials showing the four different objects not used in the subsequent experiment (spoon, mug, sickle, birdhouse). During the experiment, participants were asked to fixate on a cross at the center of the screen and respond as quickly and accurately as they could. The experiment consisted of four blocks, each containing 96 trials, resulting in a total of 384 trials. Stimulus pairings (including exemplars) were counterbalanced across blocks, and trial order within blocks was randomized. Participants rested between blocks and initiated each block at their own pace. The experiment lasted approximately 40 minutes.

After completing the attentional blink task, participants completed a two-part co-occurrence questionnaire to confirm that the objects were correctly assigned to their respective contexts. The first part tested object-scene co-occurrence by asking participants to identify each object’s typical setting (kitchen or garden). The second part measured object-object co-occurrence, with participants rating how often each object pair appears together in everyday life on a scale of 0 to 100. Results from both sections verified that the objects were strongly linked to their intended contexts. Additionally, objects from different contexts received much lower co-occurrence scores than objects from the same context (Between = 10.16, Within = 54.19, *t*(39) = 16.10, *p* < .001).

### Behavioral Analyses

Behavioral analyses were conducted using JASP (Version 0.19). T1 identification accuracy was analyzed using a three-way repeated-measures ANOVA (Lag: two vs. eight x Context: between vs. within x Category: between vs. within). The analysis of T2 detection accuracy included only trials with correct T1 responses and was evaluated using separate two-way repeated-measures ANOVAs (Context: between vs. within x Category: between vs. within) for each Lag.

## Results

### T1 Identification Accuracy

T1 object identification performance was analyzed using a 2 (Lag: two, eight) X 2 (Context: between, within) X 2 (Category: between, within) repeated-measures analysis of variance (ANOVA). The results revealed three significant main effects (Figure 2A). First, T1 performance was higher when T2 was presented at Lag 8 (M =.92, SE=.02) compared to Lag 2 (M =.90, SE=.02), *F* (1,39) = 6.04, p =.02, 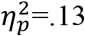. In addition, a significant contextual effect was observed, with better T1 performance when T2 belonged to a different context (M =.914, SE=.015) than when it belonged to the same context (M =.906, SE=.014), *F* (1,39) = 4.135, p = .049, 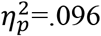. A similar effect was observed for category, with better T1 performance when T2 belonged to a different category (M =.914, SE=.014) than when it belonged to the same category (M =.906, SE=.015), F (1,39) = 4.538, p =.040,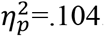. No significant interactions were observed, all *F*(1, 39) <1.

**Figure 2.**
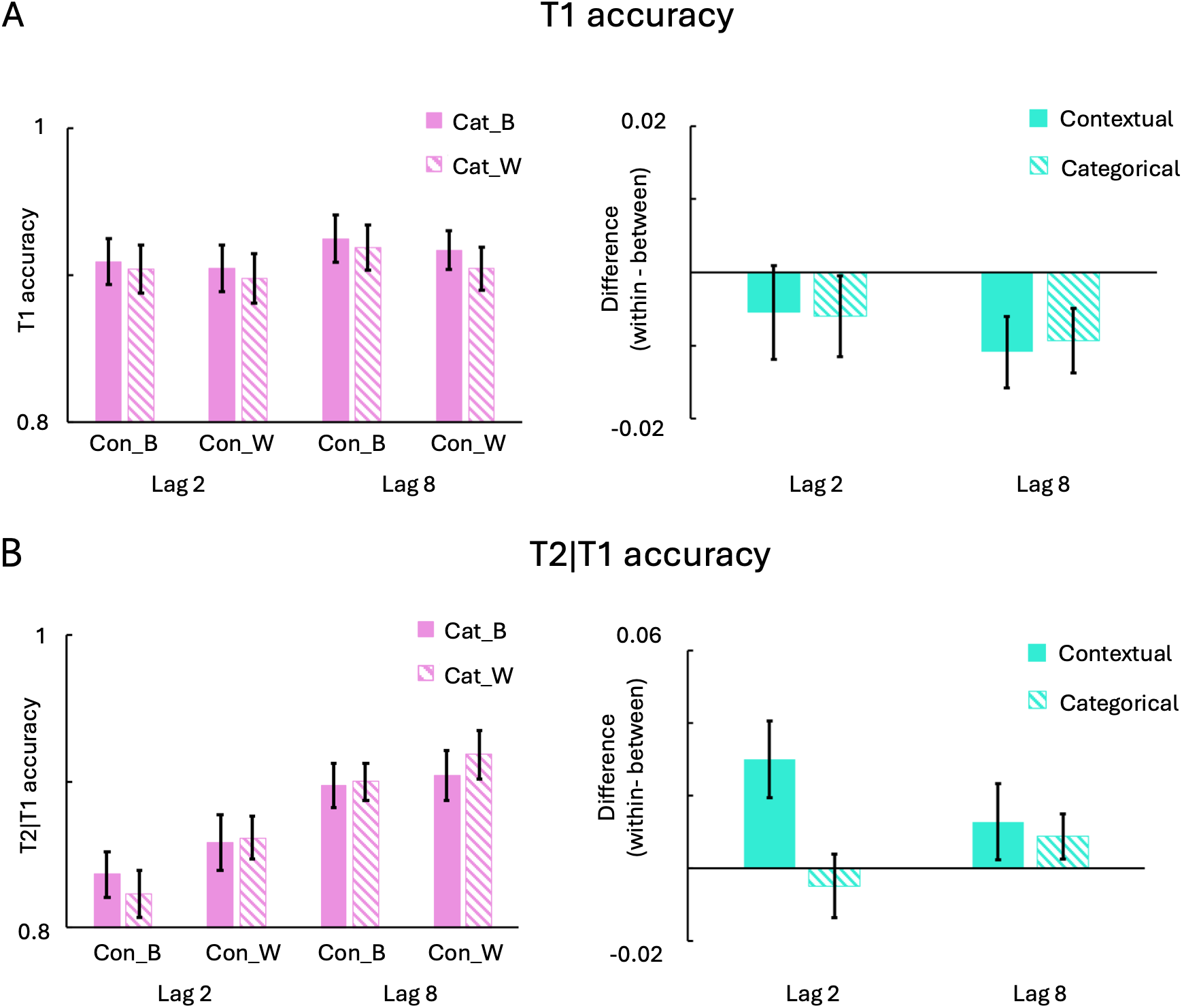
Results. (A) T1 identification accuracy. The left panel displays the accuracy for each condition. The right panel depicts the main effects of contextual and categorical relationships (within-minus between-condition accuracy), where a negative difference indicates an interference effect. (B) T2 detection accuracy (for correct T1 responses). The left panel displays the accuracy for each condition. The right panel depicts the main effects, where a positive difference indicates a facilitation effect. For both panels, error bars represent the standard error of the mean (SEM). Con_B: between-context condition; Con_W: within-context condition; Cat_B: between-category condition; Cat_W: within-category condition.

### T2 Detection Accuracy

For the T2 detection accuracy analysis, only trials with correct T1 responses were included, as the source of error is ambiguous in T1-incorrect trials (Chun & Potter, 1995). Given that the attentional blink effect is characteristically expected at shorter lags and typically resolves at longer lags, T2 accuracy was analyzed using separate 2 (Context: between, within) ×2 (Category: between, within) repeated-measures ANOVAs for the Lag 2 and Lag 8 conditions. For the Lag 2 condition, the ANOVA revealed a significant main effect of context. T2 performance was higher in the within-context condition (M =.860, SE =.016) than the between-context condition (M =.830, SE =.016), *F* (1,39) = 7.940, p =.008, 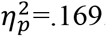. However, no significant main effect of category, *F* (1,39) = 0.325, p =.572, 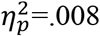, or interaction, *F* (1,39) = 1.033, p =.316, 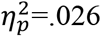, was found. The results of the two-way ANOVA for Lag 8 revealed no significant main effects: context: *F* (1,39) = 1.473, p =.232,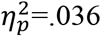; category: *F* (1,39) = 1.940, p =.172, 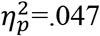, or interaction, *F* (1,39) <1.

## Discussion

By orthogonally manipulating the contextual and categorical relationships between two targets in an RSVP paradigm, we found that contextual associations attenuated the attentional blink, as indicated by improved T2 detection when the second target belonged to the same context as the first target. Furthermore, both contextual and categorical associations were associated with reduced T1 identification when the two targets belonged to the same context or category, suggesting increased interference during the maintenance of object representations in working memory.

Our results revealed a facilitative effect of contextual associations on T2 detection. Importantly, this effect was observed at Lag 2, indicating that contextual associations can modulate target processing even within the attentional blink window. These findings support the broader idea that relationships between successive events may help mitigate the attentional blink in natural environments, and further suggest that such facilitation is not limited to perceptual or categorical similarity but can also arise from contextual associations between real-world objects. Our findings are also consistent with pre-activation accounts of object processing, whereby processing of the first target activates shared representations that facilitate subsequent processing of a contextually associated second target (Lindh et al., 2019; Tang et al., 2022). This interpretation aligns with evidence that real-world objects automatically activate associated semantic representations stored in long-term memory (Bar & Aminoff, 2003; Bonner & Epstein, 2021; Yeh et al., 2026), thereby supporting efficient processing of related objects in natural environments (Collins & Loftus, 1975; Moores et al., 2003; Torralba et al., 2006; Mack & Eckstein, 2011; Yeh et al., 2025).

Interestingly, we observed an opposite pattern for T1 identification: T1 performance was reduced when the two objects shared the same context or category. Because RSVP performance depends not only on attentional selection but also on the encoding and maintenance of multiple object representations in working memory until report (Wyble et al., 2009; Martens & Wyble, 2010), this T1 impairment may reflect increased interference between overlapping object representations during post-perceptual processing. Together, these findings suggest that contextual and categorical associations can exert dissociable effects across processing stages during the attentional blink, facilitating later target processing while increasing interference at maintenance or reporting stages. However, the present data do not allow us to determine whether the T1 impairment reflects interference during maintenance or later report. Future work may apply time-resolved neural measures or tasks that separate these stages more explicitly to help clarify the mechanisms underlying the opposing effects of contextual associations on T1 and T2.

Notably, we did not observe a categorical effect on T2 detection, unlike Lindh et al. (2019). Several explanations may account for this discrepancy. First, the categories used in the present study differed from those examined in the previous study and may therefore vary in the degree of shared representations between categories (Cichy et al., 2019). Relatedly, the non-tool category in the present study was defined as a broad negative category encompassing a wide range of objects, potentially reducing representational similarity within the category. Second, in Lindh et al. (2019), shared categorical representations may have been partially driven by shared visual features, whereas perceptual similarity was systematically controlled for in the present study. Third, contextual associations may encompass richer and more specific relational information than categorical associations, so that the psychological distinction between the contextual conditions (kitchen vs. garden) may have been more salient than that between the categorical conditions (tool vs. non-tool). Consistent with this interpretation, our previous visual search study (Yeh et al., 2025) similarly found no additive effect when both contextual and categorical relationships were simultaneously available.

It is also worth noting that contextual associations in the present study were operationalized in terms of scene co-occurrence and tested using a relatively small stimulus set that was repeatedly presented across trials. The present findings, therefore, show that scene-based contextual associations are sufficient to modulate target processing during the attentional blink, but they do not yet specify which components of this contextual knowledge drive the observed facilitation. In particular, objects that co-occur within the same scene may also be linked by more specific event-based (e.g., cooking or dishwashing) or affordance-based (e.g., chopping or pouring) relationships. Future research will be needed to disentangle these forms of object-relatedness and to determine whether the observed facilitation generalizes to more variable environments.

In summary, our findings demonstrate how contextual associations shape visual cognition: they facilitate attentional access to subsequent targets while increasing interference between individuated object representations.

## Competing interest

No conflicts of interest, financial or otherwise, are declared by the authors.

## Acknowledgements

The authors thank Sirine Nouira, Pietra Pacheco Alves, and Melis Akdeniz for data collection.

## Funding

LCY is supported by the MSCA programme (101149060). DK is supported by the DFG (KA4683/5-1, project number 518483074, KA4683/7-1, project number 548389777) and an ERC Starting Grant (PEP, ERC-2022-STG 101076057). This work is further supported by the DFG under Germany’s Excellence Strategy (EXC 3066/1 “The Adaptive Mind”, project number 533717223). Views and opinions expressed are those of the authors only and do not necessarily reflect those of the funders. Neither the funders nor the granting authority can be held responsible for them.

## Data and materials availability

All study materials supporting this research are publicly available: https://osf.io/p6qyu

## Notes

https://osf.io/p6qyu

### Competing Interest Statement

The authors have declared no competing interest.

## Reference

1. Bar, M., & Aminoff, E. (2003). Cortical analysis of visual context. Neuron, 38(2), 347–358.

2. Broadbent, D. E., & Broadbent, M. H. (1987). From detection to identification: Response to multiple targets in rapid serial visual presentation. Perception & psychophysics, 42(2), 105–113.

3. Cichy, R. M., Kriegeskorte, N., Jozwik, K. M., van den Bosch, J. J., & Charest, I. (2019). The spatiotemporal neural dynamics underlying perceived similarity for real-world objects. NeuroImage, 194, 12–24.

4. Collins, A. M., & Loftus, E. F. (1975). A spreading-activation theory of semantic processing. Psychological review, 82(6), 407.

5. Deza, A., Chen, Y. C., Long, B., & Konkle, T. (2019). Accelerated texforms: alternative methods for generating unrecognizable object images with preserved mid-level features. Comput Intell Neurosci. https://github.com/ArturoDeza/Fast-Texforms.

6. Duncan, J. (1984). Selective attention and the organization of visual information. Journal of experimental psychology: General, 113(4), 501.

7. Kaiser, D., Quek, G. L., Cichy, R. M., & Peelen, M. V. (2019). Object vision in a structured world. Trends in cognitive sciences, 23(8), 672–685.

8. Kellie, F. J., & Shapiro, K. L. (2004). Object file continuity predicts attentional blink magnitude. Perception & Psychophysics, 66(4), 692–712.

9. Lindh, D., Sligte, I. G., Assecondi, S., Shapiro, K. L., & Charest, I. (2019). Conscious perception of natural images is constrained by category-related visual features. Nature communications, 10(1), 4106.

10. Long, B., Yu, C. P., & Konkle, T. (2018). Mid-level visual features underlie the high-level categorical organization of the ventral stream. Proceedings of the National Academy of Sciences, 115(38), E9015–E9024.

11. Luck, S. J., & Ford, M. A. (1998). On the role of selective attention in visual perception. Proceedings of the National Academy of Sciences, 95(3), 825–830.

12. Mack, S. C., & Eckstein, M. P. (2011). Object co-occurrence serves as a contextual cue to guide and facilitate visual search in a natural viewing environment. Journal of vision, 11(9), 9–9.

13. Martens, S., & Wyble, B. (2010). The attentional blink: Past, present, and future of a blind spot in perceptual awareness. Neuroscience & Biobehavioral Reviews, 34(6), 947–957.

14. Moores, E., Laiti, L., & Chelazzi, L. (2003). Associative knowledge controls deployment of visual selective attention. Nature neuroscience, 6(2), 182–189.

15. Olivers, C. N. (2007). The time course of attention: It is better than we thought. Current directions in psychological science, 16(1), 11–15.

16. Raymond, J. E., Shapiro, K. L., & Arnell, K. M. (1992). Temporary suppression of visual processing in an RSVP task: An attentional blink?. Journal of experimental psychology: Human perception and performance, 18(3), 849.

17. Tang, Matthew F., Kimron L. Shapiro, James T. Enns, Troy AW Visser, Jason B. Mattingley, and Ehsan Arabzadeh. “Visual awareness during the attentional blink is determined by representational similarity.” bioRxiv (2022): 2022–10.

18. Torralba, A., Oliva, A., Castelhano, M. S., & Henderson, J. M. (2006). Contextual guidance of eye movements and attention in real-world scenes: the role of global features in object search. Psychological review, 113(4), 766.

19. Weichselgartner, E., & Sperling, G. (1987). Dynamics of automatic and controlled visual attention. Science, 238(4828), 778–780.

20. Yeh, L. C., Seferovic, B., Peelen, M. V., & Kaiser, D. (2025). Contextual associations impact visual search across multiple processing stages. bioRxiv, 2025–12.

21. Yeh, L. C., Peelen, M. V., & Kaiser, D. (2026). Spatiotemporal representations of contextual associations for real-world objects. The Journal of Neuroscience, 46(19), e1967252026.

